# A genus definition for *Bacteria* and *Archaea* based on genome relatedness and taxonomic affiliation

**DOI:** 10.1101/392480

**Authors:** R.A.* Barco, G.M. Garrity, J.J. Scott, J.P. Amend, K.H. Nealson, D. Emerson

**Affiliations:** Department of Earth Sciences, University of Southern California, Los Angeles, CA, USA.; Department of Biological Sciences, University of Southern California, Los Angeles, CA, USA.; Department of Microbiology and Molecular Genetics, Michigan State University, East Lansing, MI, USA.; Smithsonian Tropical Research Institute, Panama, Republic of Panama.; Bigelow Laboratory for Ocean Sciences, East Boothbay, ME, USA.

## Abstract

Genus assignment is fundamental in the characterization of microbes, yet there is currently no unambiguous way to demarcate genera solely using standard genomic relatedness indices. Here, we propose an approach to demarcate genera that relies on the combined use of the average nucleotide identity, genome alignment fraction, and the distinction between type species and non-type species. More than 750 genomes representing type strains of species from 10 different phyla, and 19 different taxonomic orders/families in Gram-positive/negative, bacterial and archaeal lineages were tested. Overall, all 19 analyzed taxa conserved significant genomic differences between members of a genus and type species of other genera in the same taxonomic family. *Bacillus*, *Flavobacterium, Hydrogenovibrio*, *Lactococcu*s, *Methanosarcina*, *Thiomicrorhabdus*, *Thiomicrospira*, *Shewanella*, and *Vibrio* are discussed in detail. Less than 1% of the type strains analyzed need reclassification, highlighting that the adoption of the 16S rRNA gene as a taxonomic marker has provided consistency to the classification of microorganisms in recent decades. One exception to this is the genus *Bacillus* with 61% of type strains needing reclassification, including the human pathogens *B. cereus* and *B. anthracis*. The results provide a first line of evidence that the combination of genomic indices provides appropriate resolution to effectively demarcate genera within the current taxonomic framework that is based on the 16S rRNA gene. We also identify the emergence of natural breakpoints at the genome level that can further help in the circumscription of genera. Altogether, these results show that a distinct difference between distant relatives and close relatives at the genome level (i.e., genomic coherence) is an emergent property of genera in *Bacteria* and *Archaea*.

## Introduction

At the time of writing, 17,300 bacterial and archaeal species and over 3,000 genera with validly published names have been described in the taxonomic literature (Garrity, 2016; Namesforlife Database); however, based on simple phylogenetic definition there are over 200,000 bacterial and archaeal species and 60,000 genera so far detected in the SILVA database (Quast *et al*., 2013; Yarza *et al.*, 2014). Sequence data stored in the Joint Genome Institute (JGI) database, which includes data from other databases, has increased exponentially over the past decade (Markowitz *et al.*, 2015) with >130,000 genomes of bacterial and archaeal isolates, >6,000 metagenome-assembled genomes, and >2,000 single-cell amplified genomes currently available. Because the number of available datasets is constantly increasing (Mukherjee *et al*., 2017), it is becoming easier to access data that represent taxa of interest. Consequently, this added layer of available information could aid in the formal characterization of microorganisms. An essential aspect of this characterization process is the proper assignment of genus and species.

Historically, DNA-DNA hybridization (DDH) has been the ‘gold standard’ for species delineation with a DDH value of ≥70% being recognized as the species boundary between two strains (Tindall *et al.*, 2010 and other references therein). Stackebrandt and Goebel (1994) conducted a correlation analysis between DDH values and 16S rRNA gene sequence identities, and based on this, proposed a boundary of 16S rRNA gene sequence similarity of 97% for species delineation. This value, still largely used today for operational-taxonomic-unit (OTU) based analysis of microbial communities, has been updated by Stackebrandt and Ebers (2006) to a value between 98.7-99.0%, based on a greater amount of available sequence data. Subsequently, as genome sequencing has become common, whole genome comparisons became possible leading to the advent of genome relatedness indices such as the average nucleotide identity (ANI), amino acid identity (AAI), and digital DDH. More recently, Kim *et al.* (2014) proposed a 16S rRNA gene sequence similarity threshold value of 98.65% for species delineation as it equated ANI values of 95-96%, which in turn have been equated to the classical standard species delineation threshold DDH value of 70% (Goris *et al*., 2007; Richter and Rosselló-Móra, 2009). A method that only relies on protein-coding genes (i.e. neither rRNA nor tRNA are included in analysis) is the Microbial Species Identifier (MiSI), which employs both alignment fractions (AF) and ANI for demarcation of species, recommending threshold AF:ANI values of 0.6:96.5% using complete or nearly complete genomes (Varghese *et al.*, 2015).

Despite these advancements in resolving species delineation, practical guidelines that incorporate genomic properties to demarcate genera have been lacking even though genus assignment is key to performing meaningful comparisons regarding the physiology, metabolism and genomic potential of microbes. Methods to demarcate genera have been proposed that are based on either the amino acid identity (AAI) or the percentage of conserved proteins (Konstantinidis and Tiedje, 2007; Qin *et al*., 2014). However, these methods directly rely on the 16S rRNA gene sequence, which is in some cases, insensitive to evolutionary changes in the rest of the genome of a given organism, as revealed by different species sharing >99% identity over the length of this gene. Additionally, the generally-used arbitrary genus threshold of 95% 16S rRNA gene identity has been recently revisited to a lower minimum value of 94.5% with confidence interval of 94.55-95.05 and median sequence identity of 96.4% (Yarza *et al.*, 2014). In borderline cases, interpretation of results may be unclear if there are no alternative ways to confirm genus assignment. This is also the case for microorganisms with multiple, highly-divergent 16S rRNA genes. Here, we propose a novel approach that is based on the application of the MiSI method (Varghese *et al.*, 2015) and provide an objective and reproducible method of delimiting genera. We implement this approach and method by testing a variety of taxonomic groups of *Bacteria* and *Archaea*. Furthermore, supporting evidence is presented to support the reclassification of *Hydrogenovibrio halophilus* and the extensive rearrangement of species within *Bacillus,* including the human pathogens *B. anthracis* and *B. cereus*.

## Materials and Methods

### ANI and AF

ANI and alignment fraction (AF) values were obtained by the Microbial Species Identifier (MiSI) method (https://ani.jgi-psf.org/html/; Varghese *et al.*, 2015), which is also implemented in the JGI-Integrated Microbial Genomes (IMG) system (https://img.jgi.doe.gov/cgi-bin/m/main.cgi) via its Pairwise ANI tool. ANI, as defined by Varghese *et al.* (2015), is calculated for a pair of genomes by averaging the nucleotide identity of orthologous genes identified as bidirectional best hits (BBHs), which are the genes that show ≥70% sequence identity and ≥70% alignment of the shorter gene. AF, as defined by Varghese *et al.* (2015), is calculated as a fraction of the sum of the lengths of BBH genes divided by the sum of the lengths of all genes in a genome.

### Strains

The Namesforlife Database (http://www.namesforlife.com; NamesforLife, LLC, East

Lansing, MI; Garrity, 2010) was primarily used to retrieve information about type strains and type species associated with different genera. Complementary to this, equivalent strain numbers assigned by different biological resource centers (e.g., DSMZ - German Collection of Microorganism and Cell Cultures) were searched in their respective online catalogues. In addition to the above, primary literature sources were used to confirm some of these designations. All strain designations were cross-referenced in at least two databases.

### Genomes and 16S rRNA gene sequences

Genomes were obtained from IMG or the National Center for Biotechnology Information (NCBI; https://www.ncbi.nlm.nih.gov/). Generally, phyla with taxonomic families/orders that have available genomes of ≥4 type species of genera and ≥4 non-type species in a given genus were considered. Most of the genomes used in this study were sequenced directly by the JGI (47%), National Institute of Technology and Evaluation (https://www.nite.go.jp/; 5%), J. Craig Venter Institute (3%), and Broad Institute (3%). About 38% of the genomes included in this study were sequenced as part of the JGI-Genomic Encyclopedia of *Bacteria* and *Archaea*, a project that focuses on sequencing the genomes of type strains (Wu et al., 2009; Mukherjee et al., 2017). The rest of the genomes were sequenced by >140 different institutions. Sequences of the 16S rRNA gene were obtained from either IMG, NCBI, or EZBioCloud (http://www.ezbiocloud.net/). Alignments were separately performed in the SILVA Incremental Aligner version 1.2.11 (https://www.arb-silva.de/aligner/) and ClustalW via the Geneious platform (version R6; Biomatters, Auckland, New Zealand). Genetic distances were calculated in Geneious. PhyML (Guindon and Gascuel, 2003) was used via the Geneious platform to generate the maximum likelihood phylogenetic tree with the following settings: Hasegawa-Kishino-Yano (HKY85) substitution model, 1000 bootstraps, estimated transition/transversion ratio, estimated proportion of invariable sites, estimated gamma distribution, branch lengths and substitution rate optimized.

### Pangenome Analysis

Pangenome analysis of genomes from *Bacillus* and other genera in *Bacillaceae* were processed in anvi’o (version 4)(Eren et al., 2015) following the workflow for microbial pangenomics (merenlab.org/2016/11/08/pangenomics-v2/, last accessed 13 April, 2018; Delmont and Eren, 2018). In brief, we generated contig databases for each genome contig file using the command ‘anvi-gen-contigs-database’. Prodigal (Hyatt et al., 2010) was used to identify open reading frames and subsequently each database was populated with HMM profiles by comparison to several single-copy gene collections (Campbell et al., 2011; Rinke et al., 2013) using HMMER (Eddy, 2011). Once contig databases were generated for all genomes, we used ‘anvi-gen-genomes-storage’ to generate a master genome storage database to use in the pangenome analysis. We used the ncbi-blast option in ‘anvi-pan-genome’ to calculate gene similarity and MCL (van Dongen and Abreu-Goodger, 2012) for clustering under the following settings: minbit, 0.5; mcl inflation, 2; minimum occurrence, 2. Finally, to organize both gene clusters and genomes, we used Euclidian distance and Ward linkage.

### Statistical Analysis

All bidirectional, matched pair values of ANI (ANI1-2; ANI2-1) and AF (AF1-2; AF2-1) were reported as single averaged ANI and AF values in this study. The nonparametric Wilcoxon test was performed separately for each ANI and AF values to determine significant (P<0.05) differences between type- and non-type species. A discriminant multivariate analysis was performed using quadratic fitting methods, which resulted in probabilities of group assignment (i.e., type vs. non-type species). All statistical analyses were performed using the statistical software JMP Pro, version 12 (SAS Institute Inc.).

### Rationale

The nomenclatural type or "type" is defined by the International Code of Nomenclature of Prokaryotes (ICNP) as "that element of the taxon with which the name is permanently associated" (Parker et al., 2015). A type strain acts as the single reference for a given species. Each validly published species has a type strain. A type species, which is represented by its type strain, acts as the single reference for a given genus. Type strains and type species do not necessarily have to be the most representative of the species or genus, respectively. The rationale used in this study is that sister species in a given genus should be relatively similar to the type species of the cognate genus indicating high similarity at the genome level. Therefore, when a type species of a genus is compared to a non-type, sister species of the same genus, the AF:ANI values should be relatively high. In contrast to this, type species of different sister genera (i.e., in the same taxonomic order or family) should be relatively dissimilar at the genome level. This dissimilarity should be reflected in relatively low AF:ANI values.

### Guidelines

In addition to the genome(s) under consideration (e.g., of a cultured strain that needs classification), at least two genomes of type species in the same order or family are needed: a) the primary reference which is the phylogenetically closest relative that is a type species (i.e., the type species that all other microbes will be compared to), b) all other type species (≥1) in the same order or family. All available genomes of non-type, sister species that belong to the same genus of the primary reference are needed. In order for the results to be meaningful, a species must only be represented by the type strain unless its genome is not available, in which case a non-type strain of the same species that is ≥99 % identical in 16S rRNA gene sequence(s) (i.e., all copies of 16S rRNA gene in the genome) may be used. Accession numbers for these genomes should be reported.

### Interpretation

Every data point in the AF:ANI plot is calculated in relation to the primary reference which is defined above as the type strain of the type species of the specific genus to be analyzed; therefore, the plot represents genomic similarity to the reference genome. If the primary reference is compared against an identical genome (e.g., or a direct comparison against itself), an AF value of 1 and ANI value of 100 would result. If the primary reference is compared against relatively dissimilar genomes, the AF and ANI values will reflect relatively lower values. Thus, the plots that are generated effectively illustrate how distant other data points are to the primary reference in terms of AF and ANI values, or in other words, how distant other data points are to the upper-right corner (AF=1 and ANI=100).

When all comparisons are made in relation to the primary reference, a type species should always cluster with other type species in the same taxonomic order or family, reflecting relatively low AF and ANI values. A non-type, sister species should always cluster with other non-type, sister species of the same genus, reflecting relatively high AF and ANI values. The distinct clustering of type species and non-type, sister species should be conserved without any overlap. If a non-type, sister species is positioned within the cluster of type species in the same taxonomic order or family, then this indicates that the non-type, sister species could potentially be assigned to a new genus, as it is as genomically different to the primary reference as are other type species in the same taxonomic order or family. If a type species is positioned within the cluster of non-type, sister species, then there is support for potential reclassification of that species within the same genus of the primary reference, given the accumulation of other supporting evidence (e.g., 16S rRNA gene identity and phenotypic properties). It is emphasized that any proposed taxonomic rearrangement or reclassification based on AF:ANI results will necessarily affect nomenclature; therefore, any corresponding changes in nomenclature resulting from such proposal must be in alignment with the ICNP (Parker et al., 2015).

### Results and Discussion

Table 1 summarizes the 19 separate taxonomic groups that were investigated in this study along with results of statistical tests for both AF and ANI. A total of 790 genomes including 302 type species and 488 non-type species, representing 705 type strains and 10 phyla were used. For each taxonomic group, the combination of AF and ANI genomic indices resulted in distinct clustering of type and non-type species. The means of these clusters were significantly different in all tested cases with AF as a variable. In the majority of the cases (17/19), the means of these clusters were also significantly different with ANI as a variable. Discriminatory multivariate analyses, which give a sense of how separated the clusters are, supported the distinct clustering of type and non-type species in 18 out of 19 cases with 91-100% of the species correctly classified. A few representative cases will be discussed in more detail below.

**Table I.**
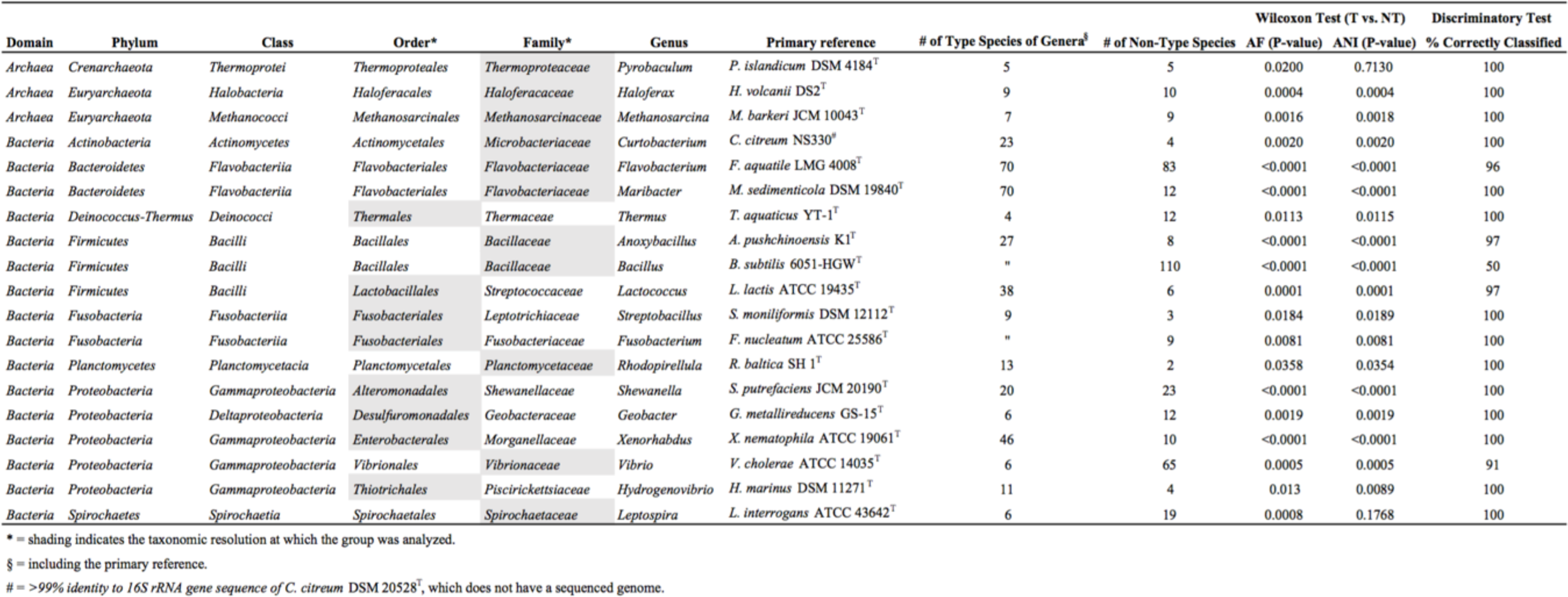
Summary of taxonomic groups and genomes included in this srudy. Gray fill-color indicates if either the taxonomic family or order was investigated. For a detailed lisl including accession numbers, see Table SI.

To initially test our approach with Gram-positive bacteria, the order *Lactobacillales* was chosen, since it contains a number of well-characterized genera and species. A continuum of AF:ANI values characterizes a typical result when intergeneric and intrageneric species of the order *Lactobacillales* (e.g., all against all) are compared without distinguishing between type and non-type strains or species (Figure 1A). Because there is no primary taxonomic reference genome to compare to, differentiation between different taxonomic groups is not possible. However, when the type species of *Lactococcus, L. lactis* subsp. *lactis* ATCC 19435^T^, is used as the primary reference, a clear distinction can be made between type species and non-type species (Figure 1B). The type species of genera in *Lactobacillales* form a distinct cluster toward lower AF:ANI values while the non-type species of *Lactococcus* cluster toward higher AF:ANI values (Table 1; Figure 1C-D). There is no overlap between these two clusters. Discriminatory multivariate analysis supported the distinct clustering of type and non-type species successfully classifying 98% of the samples.

**Figure 1.**
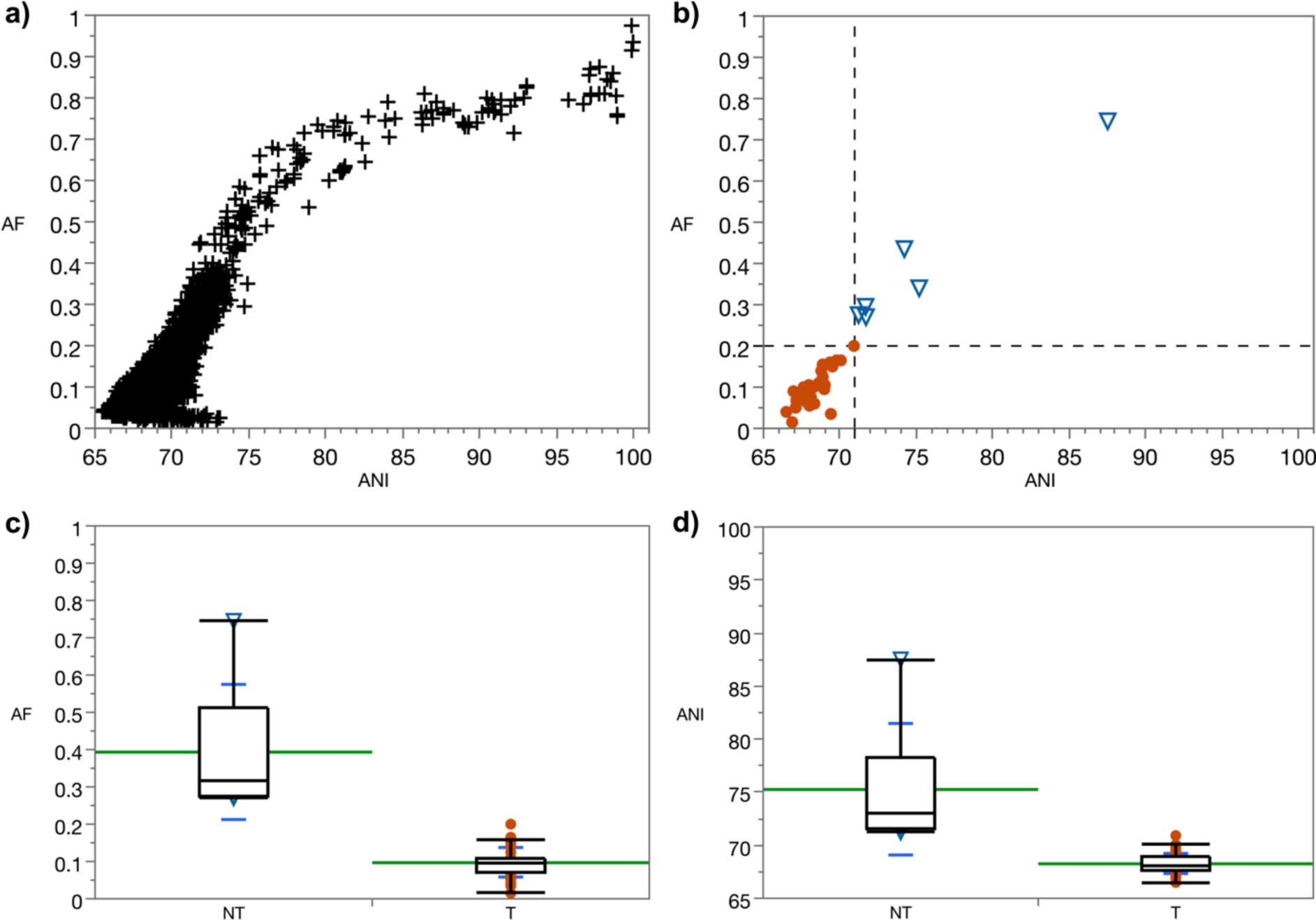
AF:ANI pairwise genome comparisons in the order *Lactobacillales*. A. Type and non-type strains/species of 15 different genera in the order *Lactobacillales* were pairwise-compared (n=6090 comparisons). B. Type species (n=37) of genera within the order *Lactobacillales* (circles) and non-type species (n=6) of the genus *Lactococcus* (in order *Lactobacillales*; triangles) were pairwise-compared *only* to the primary reference, type species *Lactococcus lactis* subsp. *lactis* ATCC 19435^T^. The bottom-left quadrant demarcates the boundary between type and non-type species. Boxplot diagrams of AF (C) and ANI (D) values as related to non-type species (NT) of the genus *Lactococcus* and type species (T) of genera within the order *Lactobacillales*. Means are shown in green. Standard deviations are shown in blue.

To test our approach with Gram-negative microorganisms, type species of genera in the order *Alteromonadales* and non-type *Shewanella* spp. were pairwise-compared to the type species *S. putrefaciens* JCM 20190^T^, which is the primary reference (Figure 2A; Table 1). A distinct clustering is seen between the type species and non-type species. Discriminatory multivariate analysis also supported the distinct clustering of type and non-type species successfully classifying 100% of the samples.

**Figure 2.**
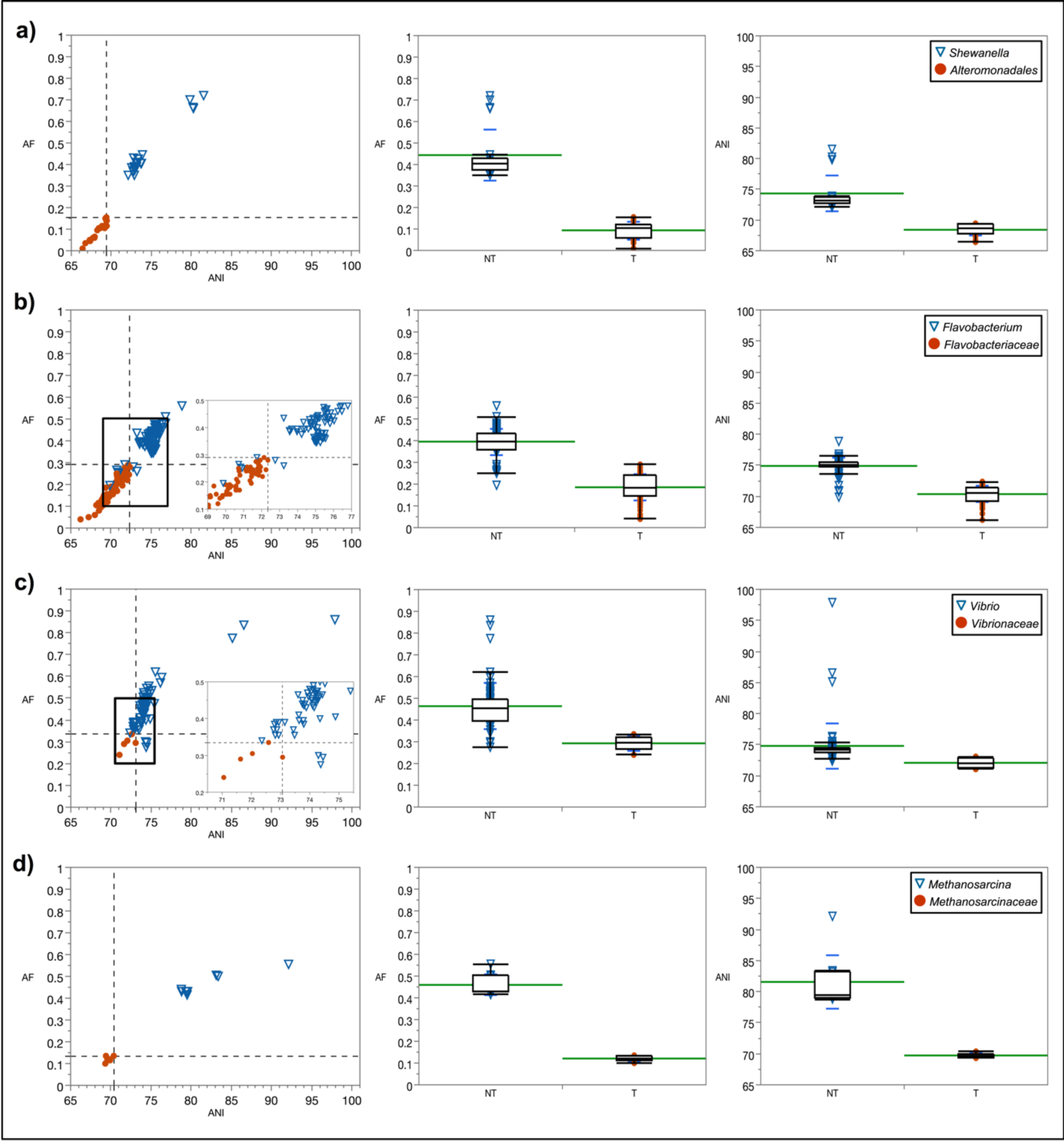
Pairwise genome comparisons to the following primary references: a) type species of the genus *Shewanella*: *S. putrefaciens* JCM 20190^T^. In circles: type species (n=19) of genera within the order *Alteromonadales*. In triangles: non-type species (n=23) of the genus *Shewanella*; b) type species of the genus *Flavobacterium*: *F. aquatile* LMG 4008^T^. In circles: type species (n=69) of genera within the family *Flavobacteriaceae*. In triangles: non-type species (n=83) of the genus *Flavobacterium*. Inset: zoomed-in boxed area; c) type species of the genus *Vibrio*: *V. cholerae* ATCC 14035^T^. Circles: type species (n=5) of genera within the family *Vibrionaceae*. Triangles: non-type *Vibrio* spp. (n=65). Inset: zoomed-in boxed area; d) type species of the genus *Methanosarcina*: *M. barkeri* JCM 10043^T^. Circles: type species (n=6) of genera within the family *Methanosarcinaceae*. Triangles: non-type *Methanosarcina* spp. (n=9). In all cases, the bottom-left quadrant demarcates the boundary between type and non-type species. Boxplots of AF (center panel) and ANI (right panel) values as related to non-type (NT) species of genus analyzed and type species (T) of genera within the order or family analyzed. Means are shown in green. Standard deviations are shown in blue.

Additional examples show that this trend is also seen with taxonomic families (Figure 2B; Table 1). The *Flavobacteriaceae* was investigated as an example of a diverse family with one of the largest number of validly published genera (currently >160). Despite this, the separate clustering of type species and non-type species was conserved. When the type species of the genus *Flavobacterium, F. aquatile* LMG 4008^T^ is used as the primary reference, all type species (i.e. with available genomes) in the *Flavobacteriaceae* clustered towards lower AF:ANI values, while the vast majority of the non-type species of *Flavobacterium* clustered toward higher AF:ANI values. Discriminatory multivariate analysis successfully classified 96% of the samples. The five non-type *Flavobacterium* spp. that were outside the AF:ANI genus boundary had 16S rRNA gene sequence identities of <94% to the primary reference. In a separate example, the type species *Vibrio cholerae* ATCC 14035^T^ was used as the primary reference and compared to non-type species in the genus *Vibrio* and other type species of genera in the family *Vibrionaceae* (Figure 2C; Table 1). A distinct clustering, without overlap, is seen between the type species and non-type species. Discriminatory multivariate analysis also supported the distinct clustering of type and non-type species with >91% of the samples successfully classified, even though the clusters were in much closer proximity to each other than in previous examples.

This approach was also tested on *Methanosarcinaceae*, an archaeal taxonomic family in the phylum *Euryarchaeota* (Figure 2D; Table 1). The primary reference was the type species *Methanosarcina barkeri* JCM 10043^T^. The results were consistent with previous examples on bacterial taxonomic groups displaying clustering of non-type species separately from type species. Discriminatory multivariate analysis successfully classified 100% of the samples. Similar results were obtained in the family *Haloferacaceae*. Additionally, the family *Thermoproteaceae* in the phylum *Crenarchaeota* was tested with *Pyrobaculum islandicum* DSM 4184^T^ as a primary reference (Figure S1). A significant distinction between type and non-type species was seen with AF (P=0.0200) but not with ANI (P=0.713). Despite this, there was no overlap of the type/non-type clusters and discriminatory multivariate analysis successfully classified 100% of the samples.

In a similar manner, genus assignment in the recently rearranged *Hydrogenovibrio-Thioalkalimicrobium-Thiomicrospira* cluster was tested as an example of a bacterial taxonomic group with historical taxonomic issues (Takai *et al*., 2004; Tourova *et al*., 2006; Figure S2). Boden *et al.* (2017) provided a detailed evaluation of the taxonomy of this cluster which belongs to the family *Piscirickettsiaceae* and proposed to place many of the species into different genera. We tested these newly proposed assignments using the approach described above and the MiSI method. When testing genus assignment to *Hydrogenovibrio*, the type species *H. marinus* DSM 11271^T^ was used as a primary reference (Figure 3). The single data point that crossed the boundaries set by the AF:ANI values between type species of genera in the family *Piscirickettsiaceae* and the non-type species of the genus *Hydrogenovibrio* belonged to *Hydrogenovibrio halophilus* DSM 15072^T^. Aside from this data point, there was no overlap of AF:ANI values between the type- and non-type species, indicating support for the assignment of *H. crunogenus*, *H. thermophilus*, and *H. kuenenii* to the genus *Hydrogenovibrio*. Similarly, the recently proposed new genus *Thiomicrorhabdus* (Boden *et al*., 2017) was also tested (Figure S3). The results indicate that the new classification is supported by AF and ANI values as the type species of genera in the family *Piscirickettsiaceae* and *Thiomicrorhabdus* spp. remain well separated. The reclassification of *Thioalkalimicrobium* spp. as *Thiomicrospira* spp. (Boden *et al*., 2017) is also supported by ANI and AF values (Figure S4), as the *Thioalkalimicrobium* spp. (now non-type *Thiomicrospira* spp.) cluster away from the type species of genera in the family *Piscirickettsiaceae*.

**Figure 3.**
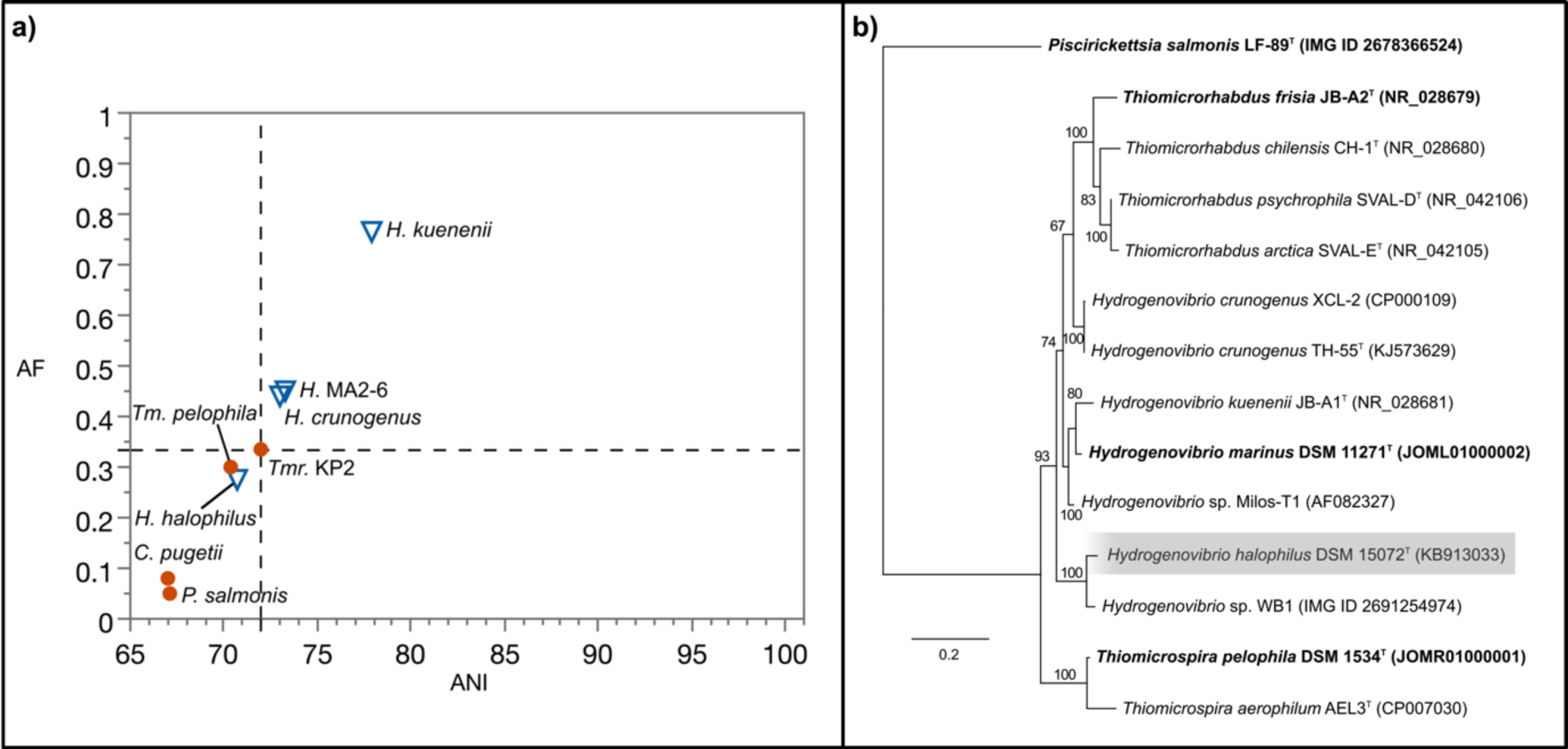
Analysis of *Hydrogenovibrio*, as rearranged by Boden *et al.* (2007). A. Pairwise genome comparisons to the primary reference *Hydrogenovibrio marinus* DSM 11271^T^. Non-type species within the genus *Hydrogenovibrio* are shown in triangles. The type strain *H. thermophilus* I78^T^ does not have a sequenced genome; therefore, *H.* sp. MA2-6 (>99% pairwise identity in 16S rRNA gene sequence) was used instead. Similarly, *H. crunogenus* XCL-2 (>99% pairwise identity in 16S rRNA gene sequence) is used instead of the type strain *H. crunogenus* TH-55^T^, as the latter does not have a sequenced genome. Type species of genera within the family *Piscirickettsiaceae* are shown in circles. The bottom-left quadrant demarcates the boundary between type and non-type species. B. Maximum-likelihood phylogenetic tree based on an alignment of 16S rRNA genes indicating the phylogenetic positioning of *H. halophilus* DSM 15072^T^. Type species of genera are shown in bold. The scale bar indicates 20% sequence divergence.

Our approach has identified a potential misclassification of *H. halophilus* (basonym *Thiomicrospira halophila*) because it clusters with the type species of genera in the family *Piscirickettsiaceae,* other than *Hydrogenovibrio*. Below we show additional evidence that could support a potential reclassification of *H. halophilus*. A closer look at the multiple 16S rRNA gene sequences of *H. halophilus* revealed that they have 94.0-94.2% pairwise-identities to the type species, *H. marinus* DSM 11271^T^. Our results differ from those of Boden *et al.*, who reported 95.7 % sequence identity (2017) between *H. halophilus* and *H. marinus*. The difference in identity appears to be due to differences in the 16S rRNA gene sequence between the near-full-length version of Sorokin *et al*. (2006; 1420 bp; accession number DQ390450) and the full-length versions of this gene (1439 bp each; IMG ID 2518265101 and 2518265550) originating from a draft genome (Scott *et al.,* 2018) and used in this study. The phylogenetic placement of this organism also supports the idea that *H. halophilus* is distinctly positioned in its own clade as it branches away from the cluster of other *Hydrogenovibrio* spp. (Figure 3B). One differentiating characteristic of *H. halophilus* is the DNA G+C content of 56.6%, much higher than the ca. 44% of the other newly classified *Hydrogenovibrio* spp. Another major differentiating aspect is the higher NaCl optimum/maximum of 1.5/3.5 M for *H. halophilus* versus the lower 0.2-0.5/0.6-1.2 M for other *Hydrogenovibrio* spp. Therefore, based on AF and ANI results, molecular, phylogenetic, and physiological evidence, *H. halophilus* could potentially be assigned to a new genus.

The *Bacillaceae* was also investigated as an example of taxonomic family that is medically relevant (Figure 4A). Generally, there were significant differences between non-type *Bacillus* species and type species of genera in the family *Bacillaceae* when *B. subtilis* 6051^T^ (Kabisch et al., 2013) was used as a primary reference (Table 1; Figure S5). However, a large number of non-type *Bacillus* species clustered with type species (50% misclassified by multivariate, discriminatory method), which strongly suggests that the genus *Bacillus* is in need of an extensive and thorough rearrangement. Of note, it is highlighted that human pathogens *B. anthracis* and *B. cereus* cluster with type species of genera in *Bacillaceae*, indicating that they are at least as genomically different to *B. subtilis* as are other type species in this family, warranting a taxonomic rearrangement. As shown above with *H. halophilus*, a similar case could be made for the taxonomic rearrangement of each one of these species (e.g., <94.5% identity in 16S rRNA gene sequence to the type species *B. subtilis* 6051^T^). Compared to our analyses of other taxonomic groups, these results are atypical in the sense that the non-type species cluster overlaps considerably with the type species cluster. We highlight the fact that a large number of these specimens were designated non-type species of *Bacillus* prior to the advent of the universal use of 16S rRNA gene as a taxonomic marker in the 1990s (i.e., >20% of all type strains of non-type *Bacillus* with available genomes in our dataset). For example, *B. anthracis* was first described in 1872 (Cohn). When the primary reference is changed to *Anoxybacillus pushchinoensis* K1^T^, a type species of a genus established in 2000 (Pikuta *et al*.) using the taxonomic framework of the 16S rRNA gene, the clustering of type species of genera in *Bacillaceae* and non-type species of *Anoxybacillus* is distinct and without overlap (Table 1).

**Figure 4.**
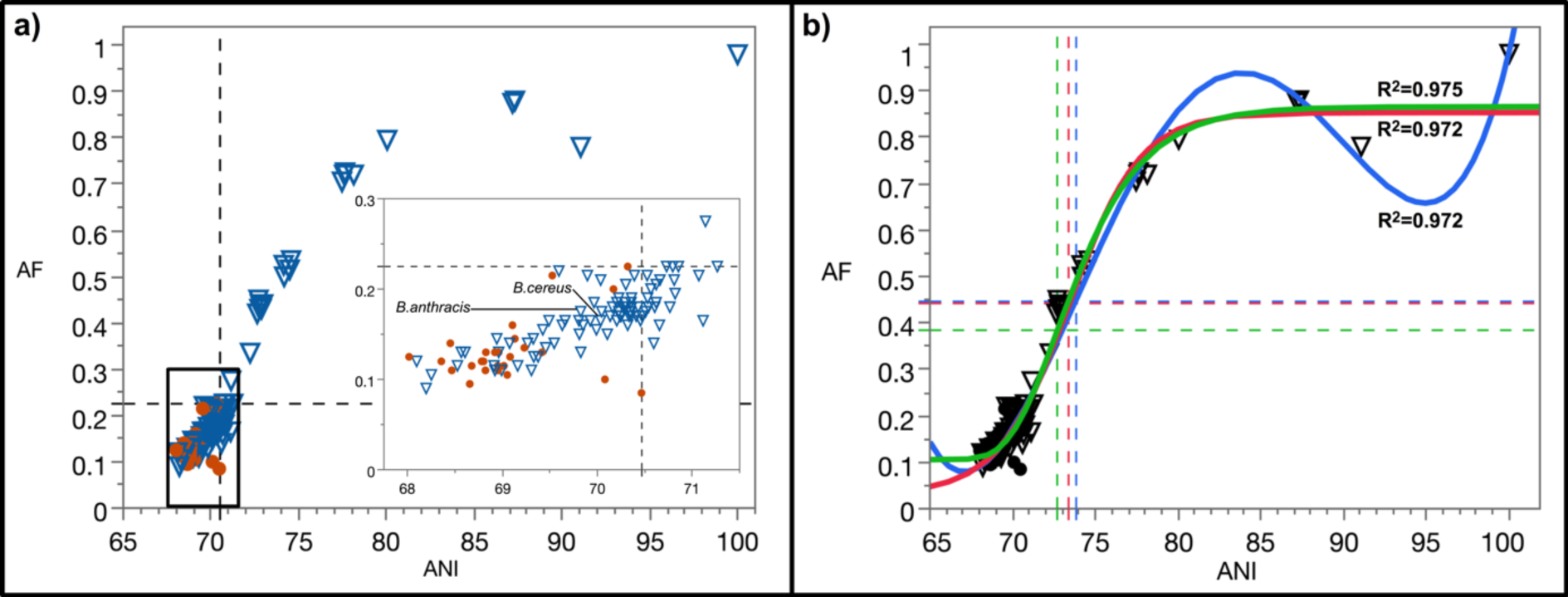
Pairwise genome comparisons to the type species of the genus *Bacillus*: *B. subtilis* ATCC 6051^T^. Circles: type species (n=26) of genera within the family *Bacillaceae*. Triangles: non-type *Bacillus* spp. (n=110). a) The bottom-left quadrant (black dashed-line) demarcates the boundary between type and non-type species; inset: zoomed-in boxed area. b) Estimated inflection points (colored dashed-lines) for *Bacillus* based on quartic (blue), logistic 4P (red), and Gompertz 4P (green) functions. The quartic function is forced to go through the AF:ANI value of 1:100.

The AF:ANI plots reveal different rates of differentiation at the DNA level. In the few cases where there are sufficient samples to analyze, and values are evenly distributed, the AF:ANI plots reveal a polynomial shape approximating a quartic function, with R^2^ values >0.97. A second derivative of this function results in a quadratic function that can be used to determine inflection points with precision. Estimates based on several examples in this study indicate two inflection points. The first inflection point at 71-74% ANI, generally approximates the genus demarcation line in all cases herein presented based on taxonomic affiliation. However, the distribution of non-type, sister species in a given genus generally includes values that fall below this inflection point at lower AF:ANI values, suggesting a potential underestimation of genera diversity within a taxonomic order or family. Estimation of this genus inflection point via Logistic and Gompertz functions generally agree with the estimations by quartic function and their use is recommended for accuracy. The second inflection point at 89-90% ANI, is consistently 5-7% lower than current ANI species thresholds (95-96.5% ANI), and is only seen with the quartic function. Because the second inflection point potentially deals with species delineation, detailed exploration (e.g., using subspecies type strains) and discussion of this topic are outside of the scope of this manuscript. However, it is noted that ANI values of 89-90% largely correspond to 16S rRNA gene identity values of ≥98.65 (Figure 3 in Kim et al., 2014).

The genus inflection point serves as practical guide for maximum AF:ANI values of genus boundary and for the identification of a genus transition zone. Ideally, and assuming abundance of type strains genomes and consequently data points, the estimated genus inflection point should closely match the genus boundary. As an example, three different functions were used to estimate the genus inflection point in *Bacillus* (Figure 4B). The obtained values give an estimation of this inflection point at AF=0.39-0.45 and ANI=72.71-73.80. An analysis of the pangenome of *Bacillus* using the most conservative estimated genus inflection point as a guide shows that only non-type *Bacillus* spp. above the genus inflection point cluster with the type species *B. subtilis* (the primary reference) while *B. anthracis* and *B. cereus* cluster with type species of other genera in *Bacillaceae* (Figure 5), corroborating, as pointed out above, that these two pathogens warrant reclassification and renaming. However, doing so would require considerable care so as not to raise objections over safety issues and a push for conservation of the current names over newly proposed names (Parker *et al.*, 2015). These results were further corroborated in phylogenomic analyses of *Bacillaceae* taking into account either 138 single copy genes or only ribosomal genes (Figures S13-S14).

**Figure 5.**
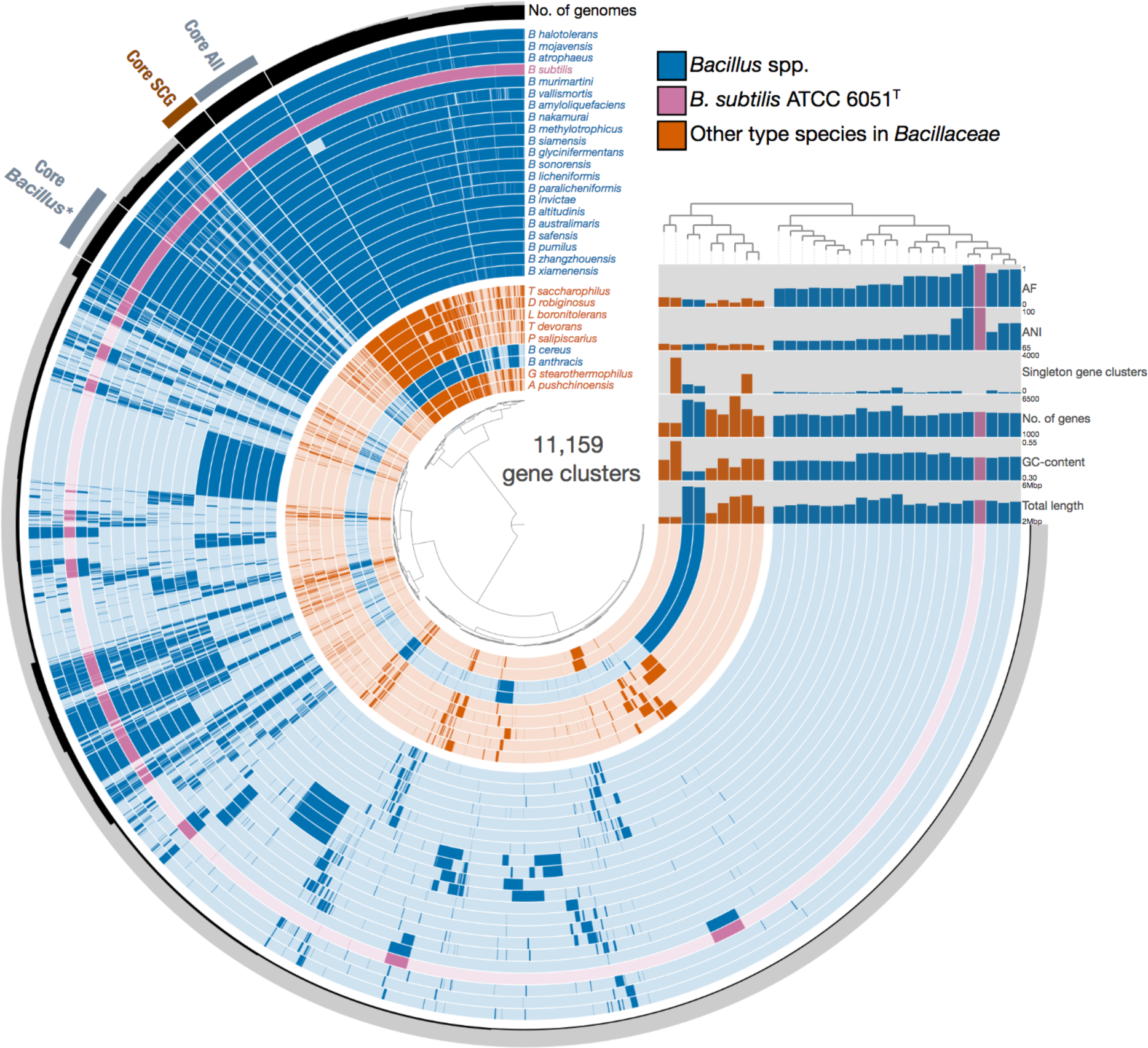
Pangenome of *Bacillus* (in blue). Type species of genera in the family *Bacillaceae* (i.e., other than *B. subtilis* ATCC 6051^T^) that are relatively close to the genus demarcation AF:ANI threshold (see Figure 4A) are shown in orange. For this pangenomic view, only *Bacillus* spp. with AF:ANI values above an estimated genus inflection point (i.e., >0.39:72.71; Figure 4B) are shown, with exception of *B. anthracis* (0.18:69.8) and *B. cereus* (0.17:70.0). ANI and AF values were obtained via pairwise genome comparisons to the primary reference, *B. subtilis* ATCC 6051^T^ (in purple). Hierarchical clustering was performed on the presence-absence of gene clusters using Euclidean distance and Ward linkage. * = *Bacillus* spp. above the estimated genus inflection point; SCG = single copy genes.

It is important to note that the genus inflection point is *estimated* based on nonlinear regression and only serves as a guide for future taxonomic designation by providing maximum AF:ANI values. This is particularly important in the case of a new taxonomic designation with a genome displaying AF:ANI values that are right above the effective genus boundary, in a transition zone; is it a novel type species or a novel non-type species? Having an estimated inflection point helps in making this decision. Thus far, this could be properly calculated only for a few taxonomic groups due to lack of genomes of type strains and/or actual isolates. *B. pumilus*, *B. safensis*, *B. invictae*, *B. altitudinis*, *B. zhangzhouensis*, *B. xiamenensis*, and *B. australimaris* are borderline species positioned within the limits of the most and least conservative estimated genus inflection points. Their 16S rRNA gene identities in relation to the primary reference range from 96.8% to 97.4%, suggesting that the genus inflection point of *Bacillus* approximates a value that is much higher than the current genus threshold based on 16S rRNA gene identity (94.5%).

The approach we have used to demarcate genera is complementary and does not replace the conventional polyphasic method of designating a genus, which includes thorough analyses of the full-length 16S rRNA gene, phylogeny, physiology and metabolism, among other aspects. The AF:ANI boundary values for genus demarcation in a given order or family can be refined as more type strains and type species are genomically sequenced and formally described. This approach emphasizes the importance of type strains and type species in the evaluation of taxonomy with genomic indices (e.g., ANI or AF), as these specimens function as reference points for their respective taxonomic groups. In this respect, our approach is in line with recommendations regarding taxonomy, including relevant comparisons to type strains and type species for the characterization of novel *Bacteria* and *Archaea* (Tindall *et al.*, 2010). It is reiterated that any rearrangement or reclassification of taxa should be in alignment with the ICNP (Parker *et al.*, 2015). In particular, a potential rearrangement of type species must be carefully analyzed as a genus can only contain one type species, and rules and principles of nomenclature must be followed in order to properly do that in accordance with the ICNP.

Uncultivated *Bacteria* and *Archaea* represented by metagenomic assembled genomes (MAGs) and/or single cell amplified genomes (SAGs) could potentially be used in the implementation of this approach; however, it is noted that type strains and type species should be used as references for such analyses, if genomes are available (see *Guidelines*). The designation of type strains and type species is governed by the ICNP and require a thorough characterization of the microorganism and the deposit of the type strain in at least two different culture collections. Naturally, the proper characterization, and therefore the type designation is, in most cases, impossible for uncultivated *Bacteria* and *Archaea* given the current guidelines set by the ICNP. This could change if genomes are subsequently considered type material, as recently proposed (Whitman, 2015) and discussed (Konstantinidis *et al*., 2017). In lieu of type strains and type species, proper and consistent genomic references must be established in order to analyze taxonomic groups without cultivated representatives. Caution is advised in the interpretation of data resulting from MAGs since they can represent composite genomes of different strains. In theory, SAGs could serve as good taxonomic frames of reference for uncultivated genera; however, genome incompleteness in SAGs is an issue that would need to be addressed.

The MiSI method explicitly excludes tRNA and rRNA genes in order to avoid inflation of AF or ANI values. However, the current taxonomic framework is largely based on the use of the 16S rRNA gene for classification. Therefore, the approach herein investigated is not completely independent of the 16S rRNA gene. The fact that in all the groups tested, with the exception of the *Bacillaceae* (note: although significant differences between type and non-type species were still detected for this group), there was strong concordance of type species of genera being delineated from non-type species of the primary reference provides evidence to the success of the use of the 16S rRNA gene within the taxonomic framework. The 16S rRNA gene has served and is continuing to serve the scientific community well, especially in maintaining consistency in classifications. However, our estimation of inflection points indicates that the 16S RNA gene identity minimum threshold value of 94.5% could underestimate genera diversity in some taxa [e.g. *Bacillaceae (Bacillus)*] and overestimate it in others [e.g. *Enterobacterales (Xenorhabdus)*], especially if multiple non-identical 16S rRNA genes are present in the genomes. It is clear that different taxonomic groups are characterized by different AF:ANI threshold values for genus demarcation. However, even within established taxonomic groups, the effective AF:ANI threshold values will still be "moving targets", dependent on the assignment and re-assignment of new type species. Thus, a single, universal, stationary threshold for genus delineation will not be able to sensitively resolve genus assignments for all taxa.

The reader is made aware that variations in bidirectional, matched pair AF values (AF1-2; AF2-1) exist. These variations are usually small, but could be relevant in the interpretation of data, especially in borderline cases. Bidirectional, matched pair ANI values (ANI1-2; ANI2-1) differ by <0.1 % in the majority of the cases, with a few cases differing by <0.3%, which would make the error bars invisible to the naked eye. Figures with corresponding standard deviation values for AF are included in a supplemental file (Figure S6-S12).

The use of ANI has been previously stated as not applicable for the demarcation of genera (Qin *et al*, 2014; Kim *et al*., 2014). Our results indicate that the MiSI method, and therefore AF and ANI, can be used to visualize natural breakpoints that can be used to circumscribe genera, if the guidelines presented in this study are followed. Adaptation of this method to demarcate higher taxonomic ranks, including phyla, has not been tested as it is beyond the scope of this study. Nonetheless, such an approach would be warranted. Finally, we highlight that the results demonstrate a conserved genomic coherence at the genus level for numerous different taxa, shedding light on a fundamental emergent property of *Bacteria* and *Archaea*, a property that has proven largely elusive at the species level.

## Acknowledgements

We would like to thank Anna Ratner at JGI for updating the interface of the Pairwise ANI tool on the IMG platform. We would also like to thank Trudy Wassenaar and Nikos Kyrpides for comments and suggestions on this manuscript. RAB was supported by the National Science Foundation postdoctoral research fellowship in Biology (award #1523639). JJS was supported by the Gordon and Betty Moore Foundation through a grant to the Smithsonian Tropical Research Institute (GBMF5603).

## Supplemental Section

**Figure S1.**
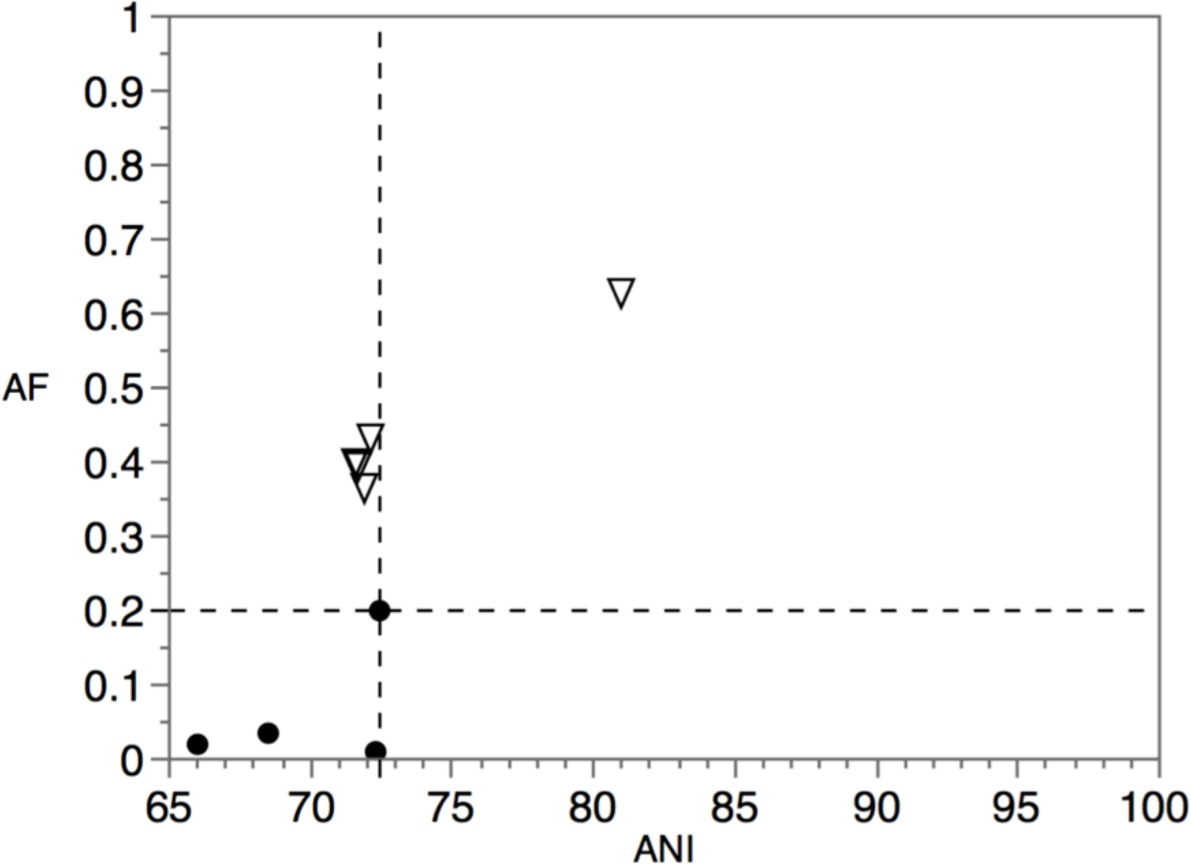
Pairwise genome comparisons to the type species of the genus *Pyrobaculum*: *P. islandicum* DSM 4184^T^. Circles: type species (n=4) of genera within the family *Thermoproteaceae*. Triangles: non-type *Pyrobaculum* spp. (n=5).

**Figure S2.**
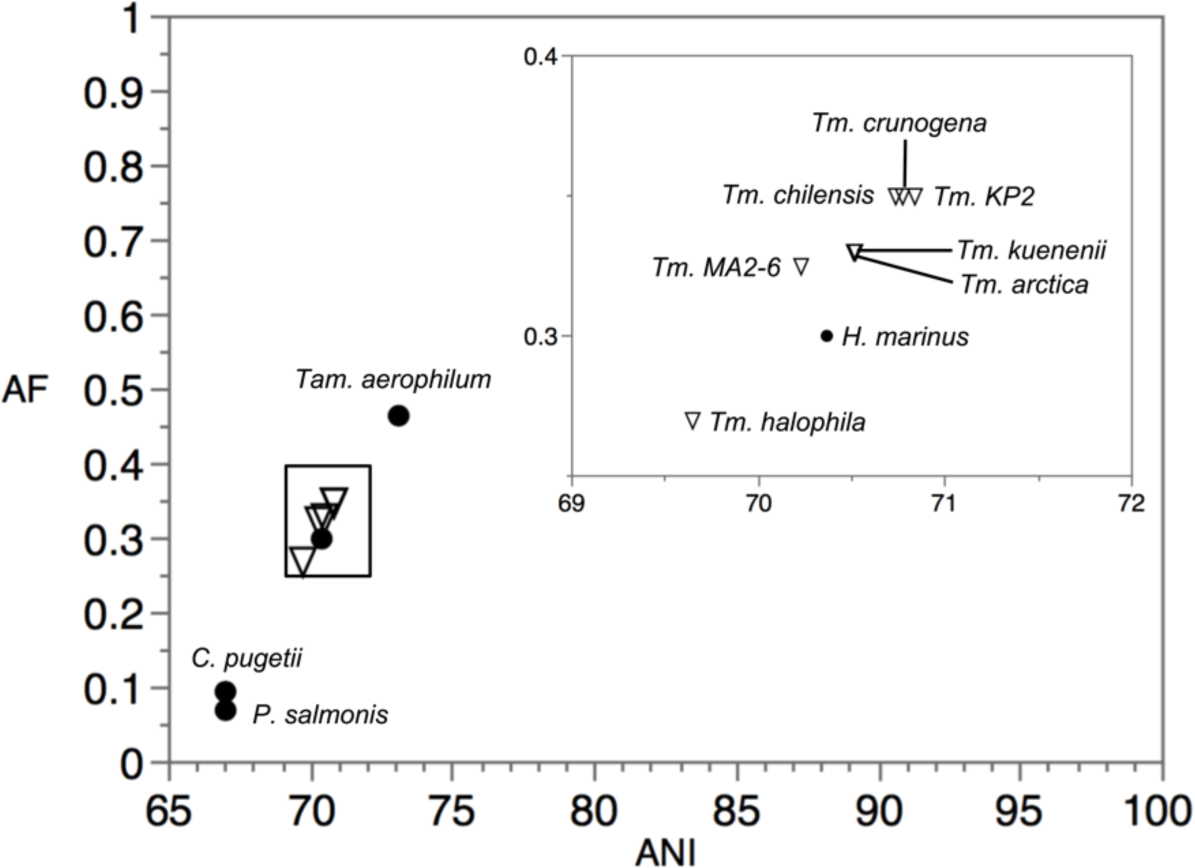
Pairwise genome comparisons to the type species *Thiomicrospira pelophia* DSM 1534^T^ using the old classification scheme (*i.e*., prior to rearrangement by Boden *et al*., 2017) show clear taxonomic issues and no distinct clustering between type and non-type species. Non-type species within the genus *Thiomicrospira* are shown in triangles. Type species of genera within the family *Piscirickettsiaceae* are shown in circles. Inset: zoomed-in boxed area.

**Figure S3.**
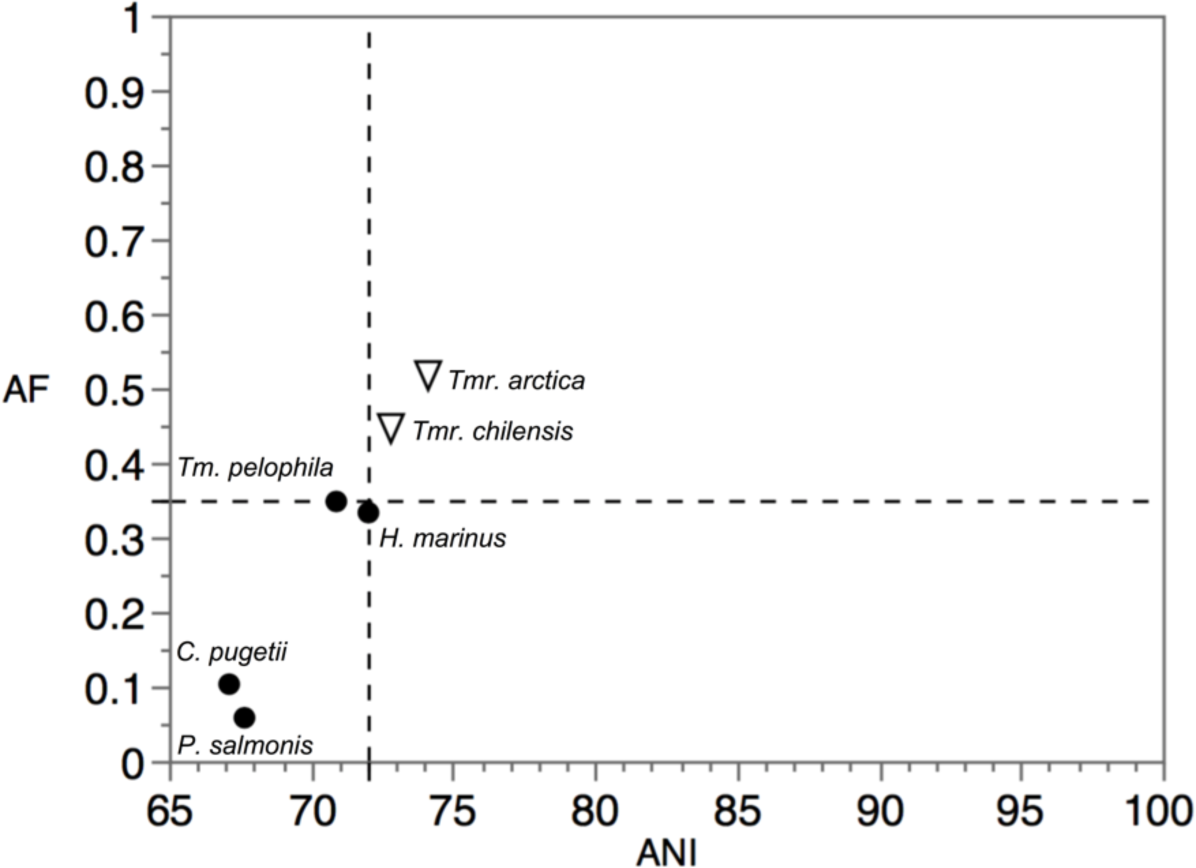
Pairwise genome comparisons to *Thiomicrorhabdus* sp. KP2 (> 99% pairwise identity to the 16S rRNA gene sequence of the type species *Tmr. frisia* JB-A2^T^, which does not have a sequenced genome) using the new re-classification scheme by Boden *et al.* (2017). Non-type species within the genus *Thiomicrorhabdus* are shown in triangles. Type species of genera within the family *Piscirickettsiaceae* are shown in circles. The bottom-left quadrant demarcates the boundary between type and non-type species.

**Figure S4.**
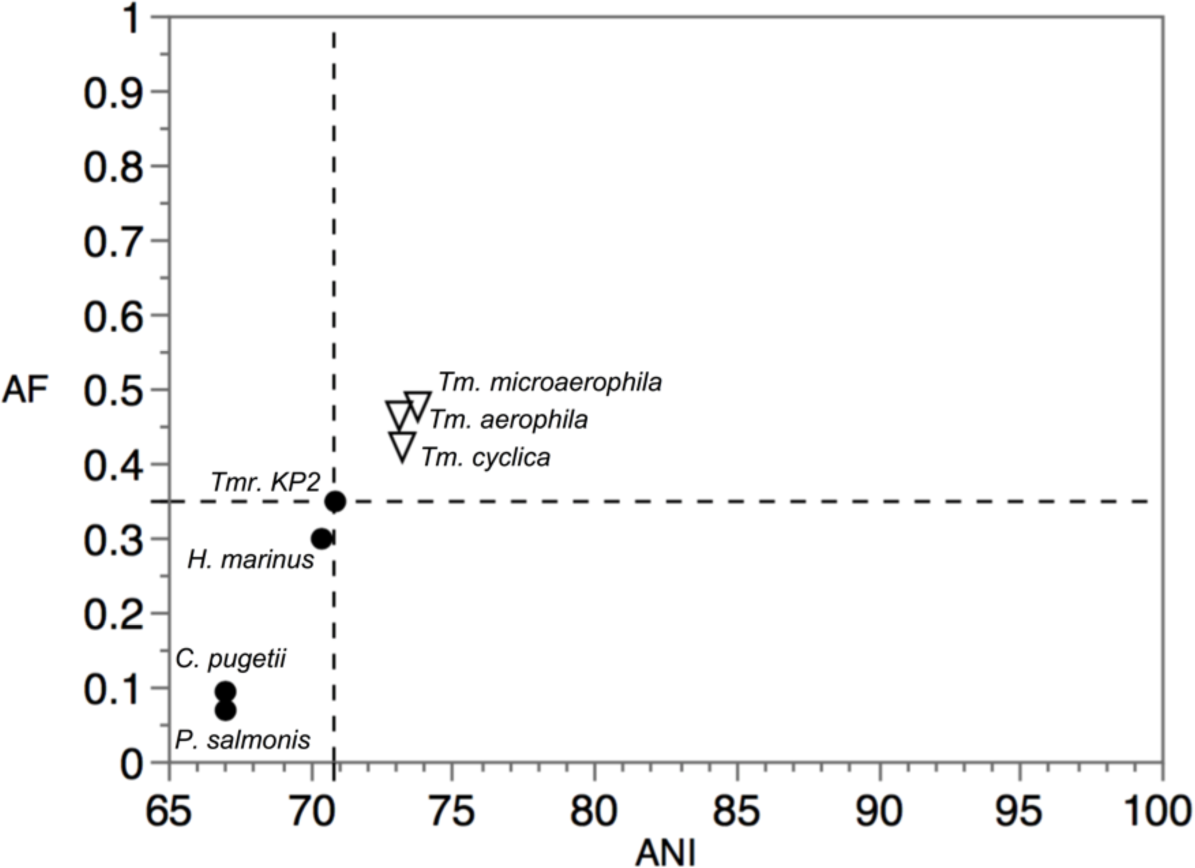
Pairwise genome comparisons to *Thiomicrospira pelophila* DSM 1534^T^ using the new re-classification scheme by Boden *et al*. (2017). Non-type species within the genus *Thiomicrospira* are shown in triangles. Type species of genera within the family *Piscirickettsiaceae* are shown in circles, with the exception of *Thiomicrorhabdus frisia* JB-A2^T^ which does not have a sequenced genome; *Thiomicrorhabdus* sp. KP2 (> 99% pairwise identity to the 16S rRNA gene sequence) was used instead. The bottom-left quadrant demarcates the boundaries between type and non-type species.

**Figure S5.**
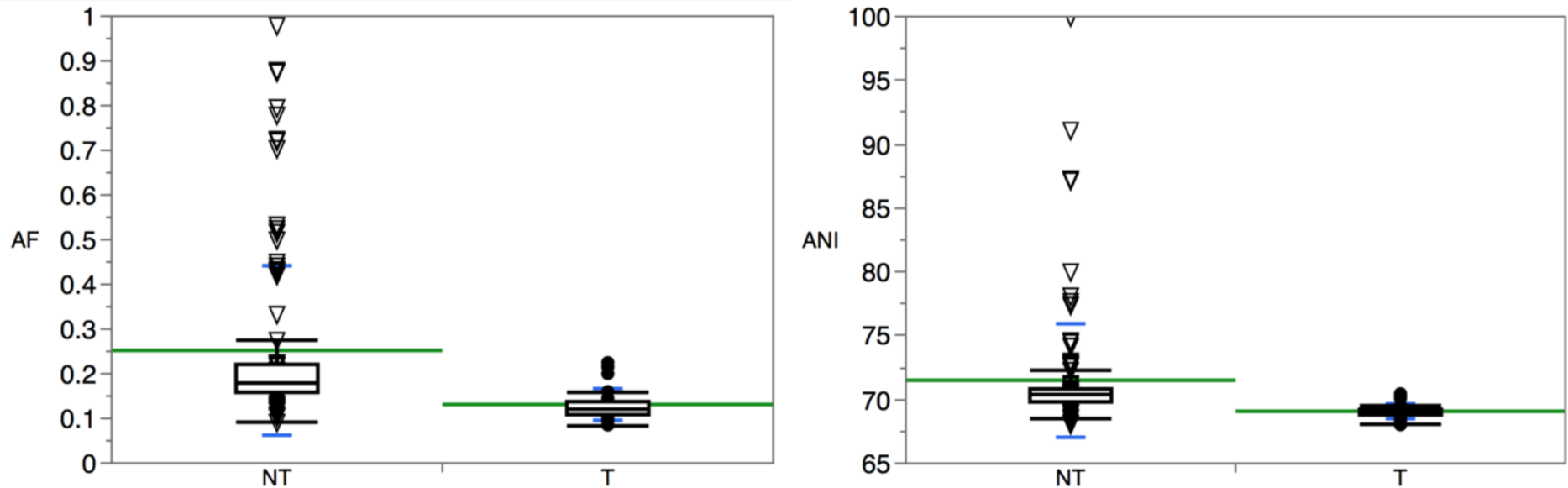
Boxplot diagrams of AF (left panel) and ANI (right panel) values as related to non-type species (NT; n=120) of the genus *Bacillus* and type species (T, n=26) of genera within the family *Bacillaceae*. Means are shown in green. Standard deviations are shown in blue.

**Figure S6.**
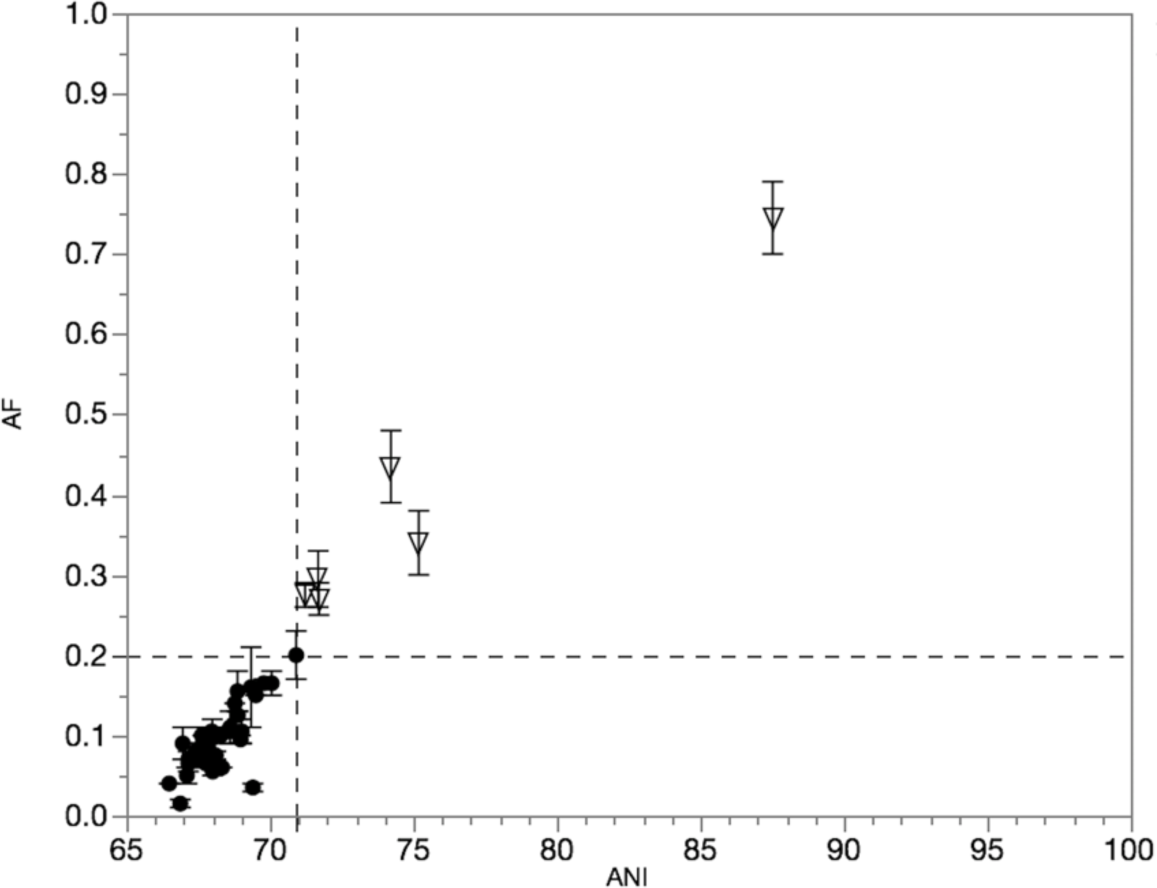
AF:ANI pairwise genome comparisons in the order *Lactobacillales* (Figure 1-right). Type species (n=37) of genera within the order *Lactobacillales* are shown in circles. Non-type species (n=6) of the genus *Lactococcus* are shown in triangles. The primary reference is the type species *Lactococcus lactis* subsp. *lactis* ATCC 19435^T^. The bottom-left quadrant demarcates the boundary between type and non-type species. Error bars indicate one standard deviation from the mean.

**Figure S7.**
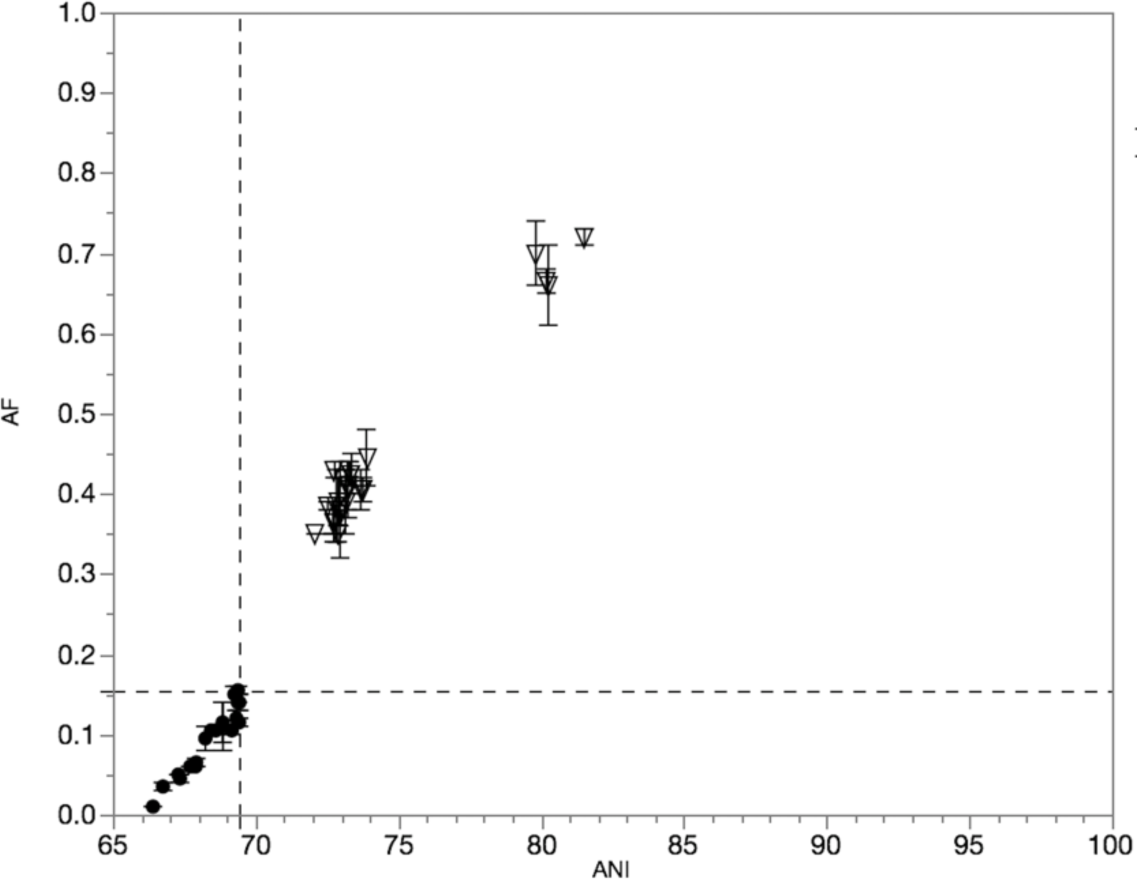
Pairwise genome comparisons to the type species of the genus *Shewanella*: *S. putrefaciens* JCM 20190^T^ (Figure 2A). In circles: type species (n=19) of genera within the order *Alteromonadales*. In triangles: non-type species (n=23) of the genus *Shewanella*. The bottom-left quadrant demarcates the boundary between type and non-type species. Error bars indicate one standard deviation from the mean.

**Figure S8.**
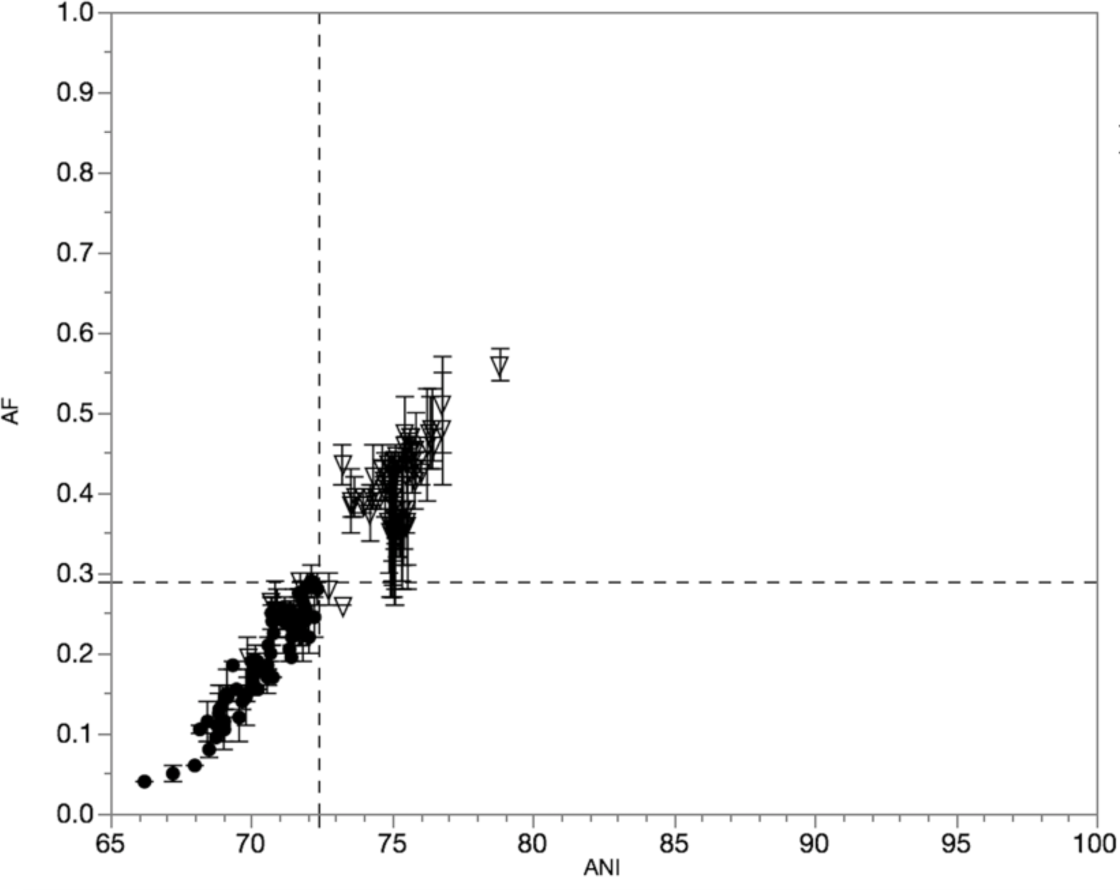
Pairwise genome comparisons to the type species of the genus *Flavobacterium*: *F. aquatile* LMG 4008^T^ (Figure 2B). In circles: type species (n=69) of genera within the family *Flavobacteriaceae*. In triangles: non-type species (n=83) of the genus *Flavobacterium*. The bottom-left quadrant demarcates the boundary between type and non-type species. Error bars indicate one standard deviation from the mean.

**Figure S9.**
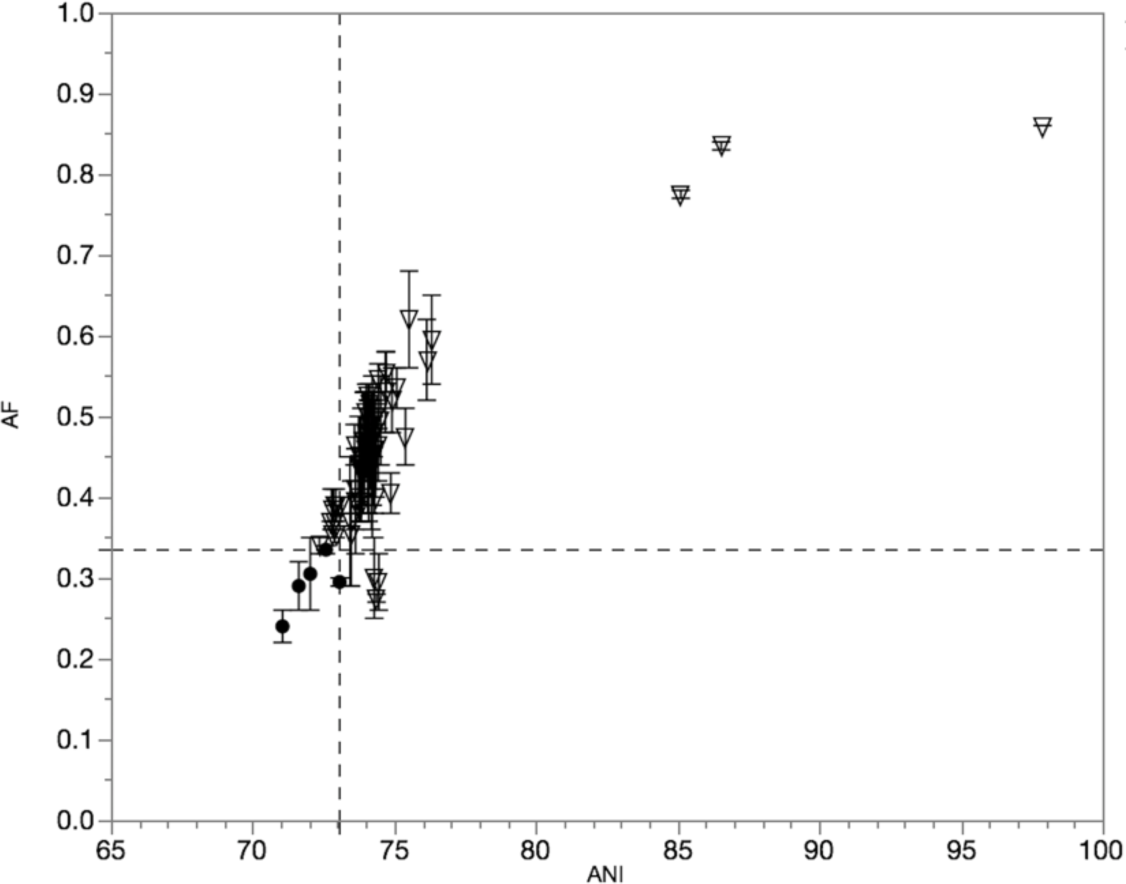
Pairwise genome comparisons to the type species of the genus *Vibrio: V. cholerae* ATCC 14035^T^ (Figure 2C). Circles: type species (n=5) of genera within the family *Vibrionaceae*. Triangles: non-type *Vibrio* spp. (n=65). The bottom-left quadrant demarcates the boundary between type and non-type species. Error bars indicate one standard deviation from the mean.

**Figure S10.**
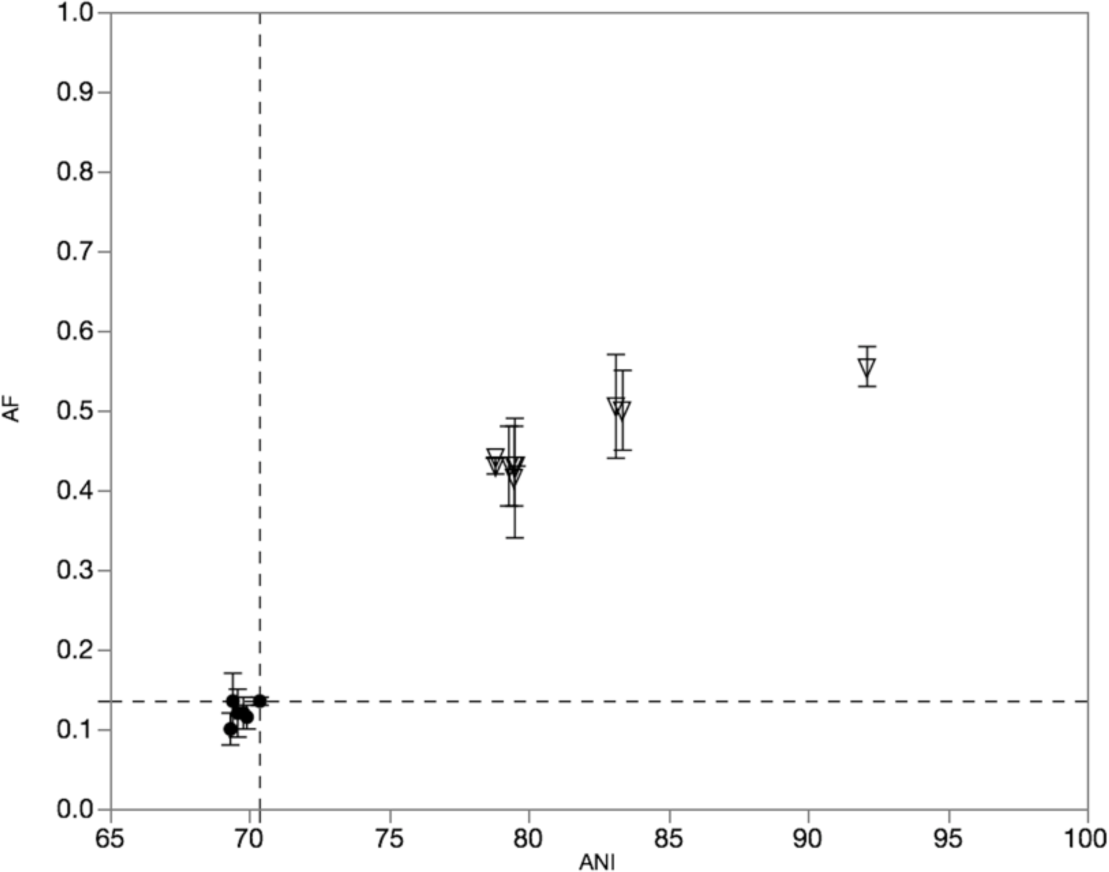
Pairwise genome comparisons to the type species of the genus *Methanosarcina*: *M. barkeri* JCM 10043^T^ (Figure 2D). Circles: type species (n=6) of genera within the family *Methanosarcinaceae*. Triangles: non-type *Methanosarcina* spp. (n=9). Error bars indicate one standard deviation from the mean.

**Figure S11.**
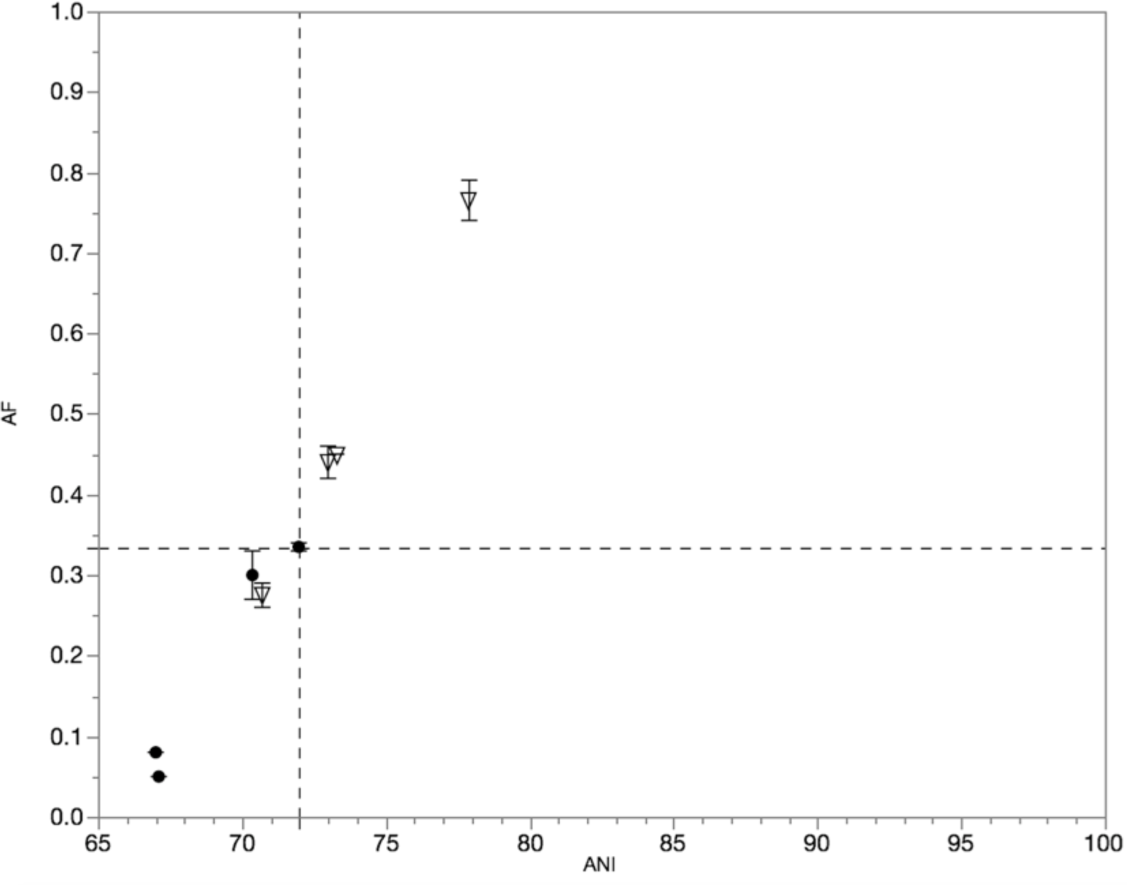
Pairwise genome comparisons to *Hydrogenovibrio marinus* DSM 11271^T^ using the new re-classification scheme by Boden *et al.* (2017)(Figure 3A). Non-type species within the genus *Hydrogenovibrio* are shown in triangles. Type species of genera within the family *Piscirickettsiaceae* are shown in circles. The bottom-left quadrant demarcates the boundary between type and non-type species. Error bars indicate one standard deviation from the mean.

**Figure S12.**
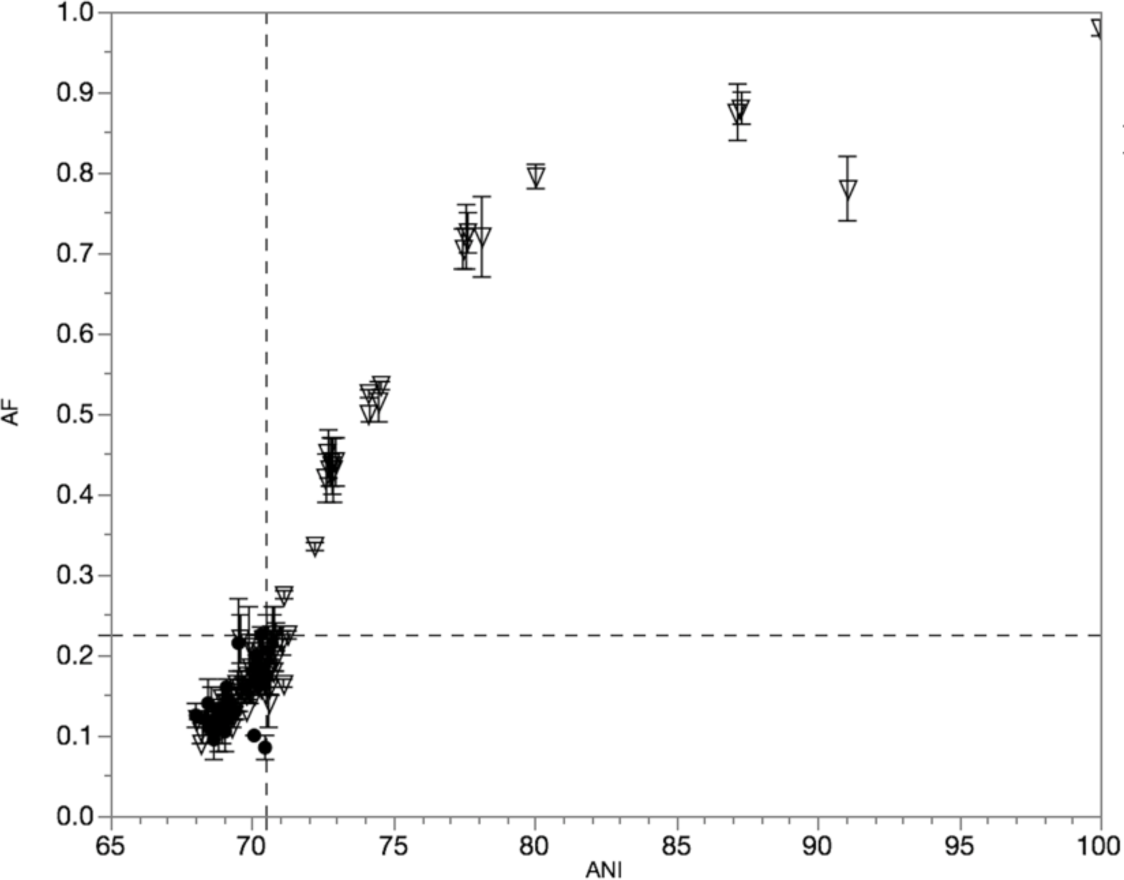
Pairwise genome comparisons to the type species of the genus *Bacillus*: *B. subtilis* ATCC 6051^T^. Circles: type species (n=26) of genera within the family *Bacillaceae*. Triangles: non-type *Bacillus* spp. (n=110). The bottom-left quadrant demarcates the boundary between type and non-type species. Error bars indicate one standard deviation from the mean.

**Figure S13.**
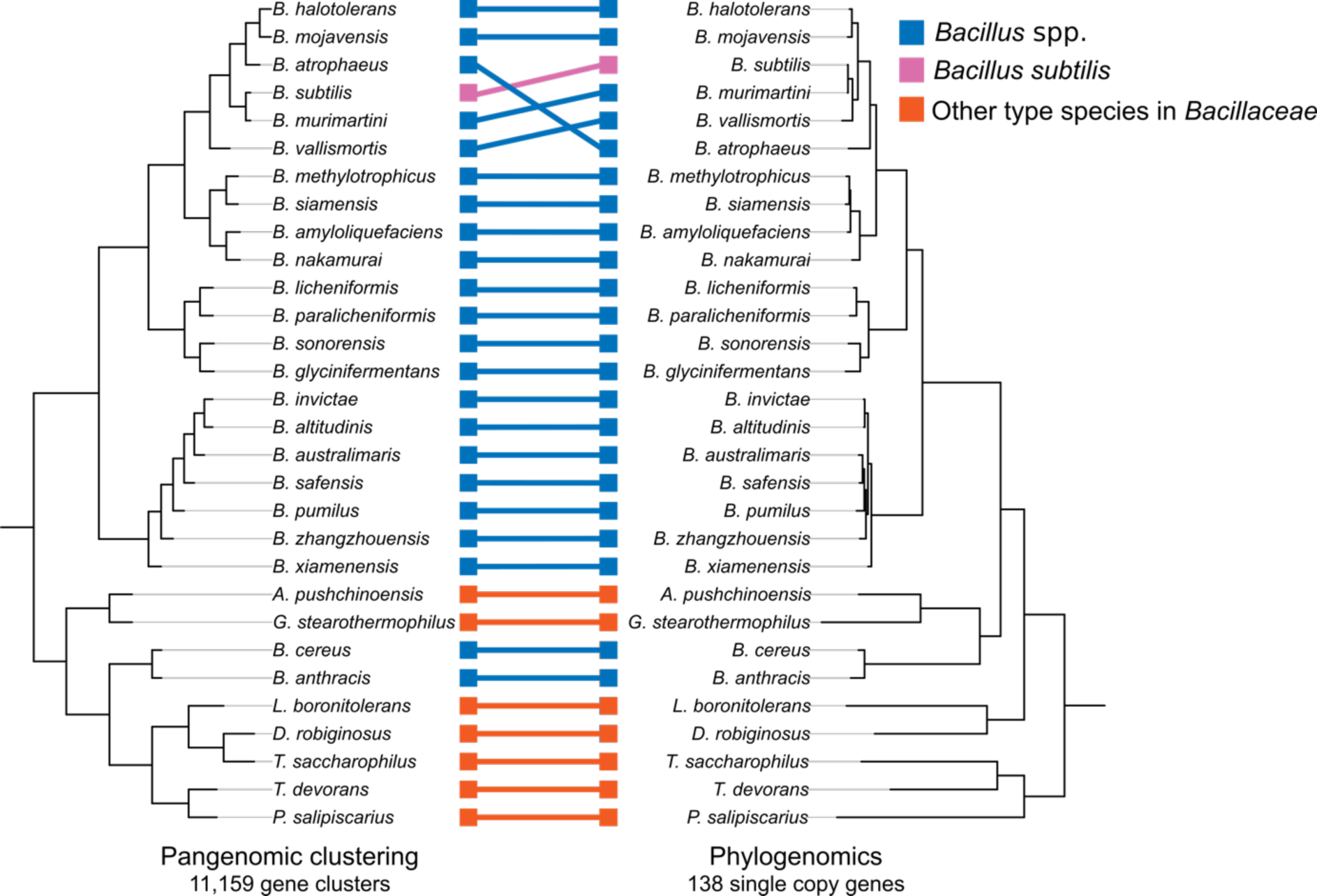
Genome clustering based on either pangenomic or phylogenomic analysis. All 138 single copy genes were included in the phylogenomic analysis.

**Figure S14.**
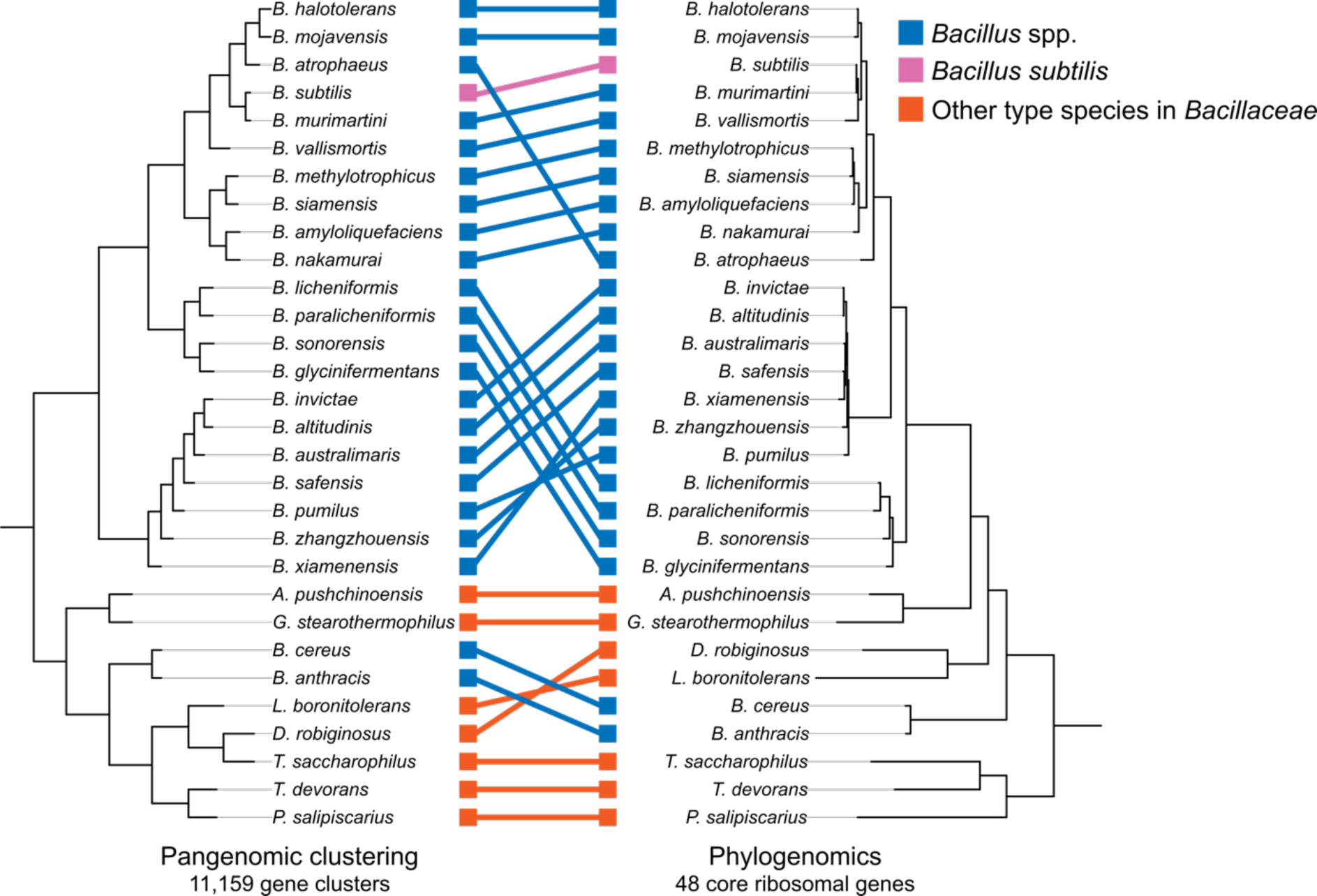
Genome clustering based on either pangenomic or phylogenomic analysis. Only the ribosomal genes were included in the phylogenomic analysis.

